# Evolution of ion channels in cetaceans: A natural experiment in the Tree of life

**DOI:** 10.1101/2023.06.15.545160

**Authors:** Cristóbal Uribe, Mariana F. Nery, Kattina Zavala, Gonzalo A. Mardones, Gonzalo Riadi, Juan C. Opazo

**Author notes:** **Corresponding authors:** Gonzalo Riadi Center for Bioinformatics, Simulation and Modeling, CBSM Department of Bioinformatics Faculty of Engineering University of Talca Talca, Chile, Juan C. Opazo Facultad de Medicina y Ciencia Universidad San Sebastián Valdivia, Chile.

## Abstract

Cetaceans could be seen as a natural experiment within the tree of life in which a mammalian lineage changed from terrestrial to aquatic habitats. This shift involved extensive phenotypic modifications, which represent an opportunity to explore the genetic bases of phenotypic diversity. Furthermore, the availability of whole genome sequences in representative species of all main cetacean groups means that we are in a golden age for such studies. Among the different molecular systems that maintain cellular homeostasis, ion channels are crucial for the proper physiological functioning of all living species. This study aims to explore the evolution of ion channels during the evolutionary history of cetaceans. To do so, we created a bioinformatic pipeline to annotate the repertoire of ion channels in the genome of the species included in our sampling. Our main results show that cetaceans have on average, fewer protein-coding genes and a higher percentage of annotated ion channels than non-cetacean mammals. Signals of positive selection were detected in ion channels related to the heart, locomotion, visual and neurological phenotypes. Interestingly, we predict that the Na_V_1.5 ion channel of most toothed whales (odontocetes) is sensitive to tetrodotoxin (TTX), similar to Na_V_1.7, given the presence of tyrosine instead of cysteine, in a specific position of the ion channel. Finally, the gene turnover rate of the cetacean crown group is more than three times faster than non-cetacean mammals.

## Introduction

Understanding the genetic basis of phenotypic diversity represents a central goal in evolutionary biology. The availability of whole-genome sequences in representative species of different groups represents a unique opportunity to advance this goal. Indeed, the Tree of Life could be seen as a set of natural experiments that help us understand different evolutionary phenomena. For example, lineages have evolved traits of biomedical interest (e.g., cancer resistance), representing an opportunity to understand how the evolutionary process solves problems from which we can potentially gain medical insights (Giroud et al. 2021; Tejada-Martinez et al. 2021; Tollis et al. 2021; Zhao et al. 2021; Oka et al. 2023; Thienel et al. 2023). Further, during the evolutionary history of vertebrates, colonizations of new habitats have resulted in severe phenotypic transformations, providing an opportunity to understand the genomic basis of diversity (McGowen et al. 2012; Nery, González, et al. 2013; Sun et al. 2013; M. Jiang et al. 2020; Bondareva et al. 2023).

The colonization of the aquatic environment by tetrapods occurred multiple times during their evolutionary history. Among mammals, the cetaceans (whales and dolphins) started transitioning from land to the sea during the Eocene around 50 Mya. A few million years later, the lineage entirely depended on the aquatic environment and diversified and occupied all the seas and many rivers of the world (Berta et al. 2005). The successful aquatic colonization from a terrestrial habitat demanded biological transformation due to gravity-related challenges, thermal regimes, a new pathogenic environment, different environmental stimuli that required sensory adaptations, and osmotic regulation, among others (Houssaye and Fish 2016; Kelley et al. 2016). Taking advantage of the advancement of genomic tools, these adaptations have been investigated from the molecular perspective to expand further our knowledge about the evolutionary process behind the conquest of an aquatic lifestyle (McGowen et al. 2011; McGowen et al. 2012; Nery, González, et al. 2013; Nery, Arroyo, et al. 2013; Sun et al. 2013; Nery et al. 2014; McGowen et al. 2020). Interestingly, several studies have reported the loss of genes as a strategy of phenotypic evolution (Feng et al. 2014; Nery et al. 2014; Sun et al. 2017; Huelsmann et al. 2019; Helsen et al. 2020; McGowen et al. 2020; Randall et al. 2022; Zheng et al. 2022; Osipova et al. 2023; Pinto et al. 2023).

Ion channels are integral membrane proteins that allow the passage of ions involved in a diverse repertoire of physiological processes. In the human and mouse genomes, 235 and 231 putative ion channels have been identified, respectively (Jegla et al. 2009). There is no doubt that ion channels represent a crucial part of the molecular machinery for the correct physiological functioning of living creatures. In fact, amino acid sequence variation has been linked to a wide range of pathological conditions, also called channelopathies (Ackerman and Clapham 1997; Ackerman 2004; Kim 2014). Thus, given their pivotal role in different physiological axes, some of which have diverged extensively in cetaceans due to the aquatic transition, it seems interesting to study their evolutionary trend in this mammalian group (Varró et al. 2021; Kashio and Tominaga 2022; Poole 2022).

This work aims to study the evolution of ion channels during the evolutionary history of cetaceans. To do so, we first created a bioinformatic pipeline to annotate the whole repertoire of ion channels in the genome of the species included in our sampling. After that, we estimated homologous relationships to study the role of positive selection and the variation in gene turnover rate. Our main results show 1) on average, cetaceans possess fewer protein-coding genes and a higher percentage of annotated ion channels than non-cetacean mammals 2) the signal of positive selection was found in ion channels related to heart, locomotion, visual and neurological phenotypes, all characteristics extensively modified in cetaceans, 3) the Na_V_1.5 ion channel of toothed whales (odontocetes), other than species of the genus *Tursiops*, is predicted to be sensitive to the potent neurotoxin tetrodotoxin (TTX), similar to Na_V_1.7, given a replacement of cysteine for a tyrosine, 4) the gene turnover rate of the cetacean crown group is more than three times faster in comparison to non-cetacean mammals.

## Material and methods

### Phylogenetic design, DNA sequences, and ion channel annotation

Our phylogenetic design included 18 mammalian species: seven toothed whales (Odontoceti) (bottlenose dolphin, *Tursiops truncatus*; orca, *Orcinus orca*; beluga, *Delphinapterus leucas*; Yangtze river dolphin, *Lipotes vexillifer*; sperm whale, *Physeter catodon, vaquita, Phocoena sinus, narwhal, Monodon monoceros*), two baleen whales (Mysticeti) (common minke whale, *Balaenoptera acutorostrata*; blue whale, *Balaenaenoptera musculus*), three artiodactyls (hippo, Hippopotamus amphibius, cow, *Bos taurus*; pig, *Sus scrofa*), one carnivore (dog, *Canis familiaris*); one perissodactyl (horse, *Equus caballus*); one chiroptera (microbat, *Myotis lucifugus*); one primate (human, *Homo sapiens*); one rodent (mouse, *Mus musculus*) and one proboscidean (African elephant, *Loxodonta africana*).

The protein-coding sequences (.faa file extension) for each species were downloaded from Ensembl v.105 (Yates et al. 2022) or the NCBI database (Sayers et al. 2022). In all cases, we kept the longest transcript for each gene. Of all the proteins present in the proteomes, we selected those that had between 2 and 35 transmembrane domains using the software TMHMM v2.0 (Krogh et al. 2001) (Fig. 1). We annotated the protein sequences based on the structural motifs using the program RPS-BLAST v2.13.0+ (with the option-outfmt 11) plus the rpsbproc package (Yang et al. 2020) (Fig. 1). We filtered the results based on the *E-value* threshold of 10^-4^ (Fig. 1). To identify the ion channels from our list of proteins, we prepared a file containing the list of ion channel conserved domains based on the Conserved Domain Database (CDD) (Wang et al. 2023). Having done that, we intersected it with the results from RPS-BLAST v2.13.0+ followed by rpsbproc. This was done using an *in-house* Perl script to identify the ion channel repertoire for all sampled species (Fig. 1). The result was the list of ion channel protein sequences from each species for the next steps (Supplementary Table S1).

**Figure 1.**
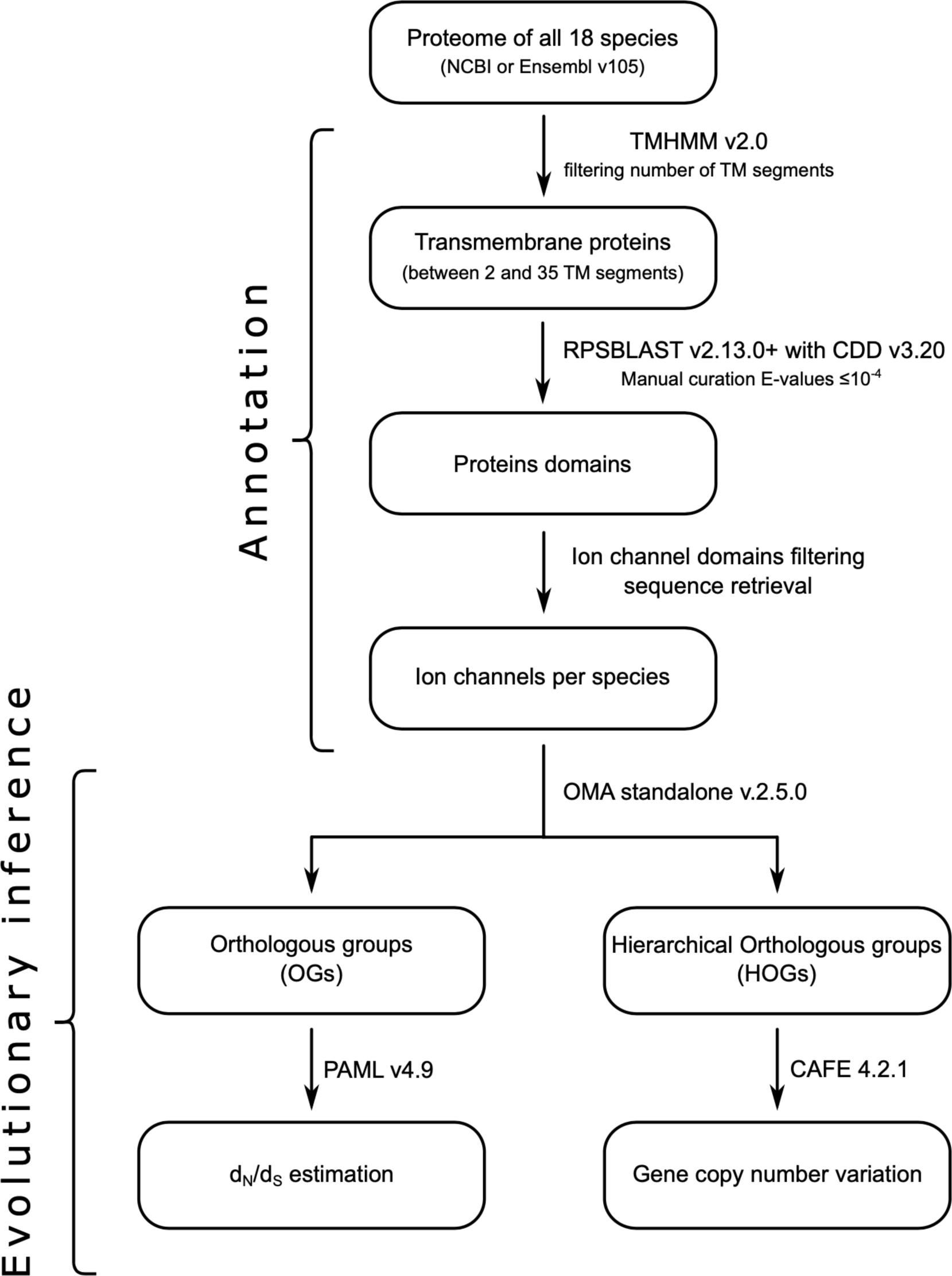
Flow diagram of the methodology used in this work. We divided the pipeline into two main steps, the annotation process, and the evolutionary inference. In the annotation process, we implemented tools like TMHMM (Krogh et al. 2001) and RPSBlast (Yang et al. 2020) to identify the repertoire of ion channels in the proteome of each species. Then, we applied the OMA standalone program to infer orthologous and hierarchical orthologous groups. After that, we used PAML (Yang 2007) to estimate the ratio of the rate of non-synonymous (dN) and synonymous substitutions (dS) (ω=dN/dS) and CAFE (Han et al. 2013) to study gene copy number variation. TM: Transmembrane

### Homology inference

After having identified the ion channel repertoire from each sampled species, the next step was to infer homologous relationships. To do this, we used the program OMA standalone v2.5 (Altenhoff et al. 2021). We inferred Orthologous Groups (OGs) containing 1:1 orthologous genes and Hierarchical Orthologous Groups (HOGs) containing the set of genes that have descended from an ancestral gene. OGs were used to perform natural selection analyses by estimating the rate of non-synonymous (d_N_) and synonymous (d_S_) substitutions. In contrast, the HOGs were used to perform gene copy number variation analyses. Both analyses take into account the phylogenetic relationships of the included species. Amino acid sequences were aligned using the FFT-NS-2 strategy of the program MAFFT v7.490 (Katoh and Standley 2013). Codon-aligned nucleotide alignments were obtained by employing the amino acid alignments as templates, using the software PAL2NAL (Suyama et al. 2006).

### Molecular evolution analyses

To test for evidence of positive selection on the cetacean orthologous groups of ion channels, we estimated the ratio of the rate of non-synonymous (d_N_) and synonymous substitutions (d_S_) (ω=d_N_/d_S_) using site models in the program PAML v4.9 (Yang 2007). For each multiple sequence alignment, we compared models that allow ω to vary among codons (M1a vs. M2a, M7 vs. M8). M2a and M8 models allow a subset of sites to have ω > 1, in contrast to the null models (M1a and M7), which only includes site classes with ω ≤ 1. All nested models were compared using the Likelihood Ratio Test (LRT), and we applied the False Discovery Rate (FDR) as a multiple-test correction (Benjamini and Hochberg 1995).

### Gene turnover rate analyses

To study variation in the gene turnover rate, we used the software CAFE v4.2.1 (Han et al. 2013). Using this program, we estimated the rate of evolution (λ) and the direction of the change regarding the size of gene families across different lineages. We implemented two models, 1) in the first model, we estimated one λ value for cetaceans as a total group (crown and stem), and a second λ for all non-cetaceans branches of the tree, and 2) in the second model, we estimated one λ for the stem Cetacea, another λ for the crown Cetacea, and a third λ for all non-cetaceans branches of the tree. The divergence time between species was obtained from the TimeTree 5 database (Kumar et al. 2022).

### Physiological phenotypes associated with the positively selected genes

To understand the physiological phenotypes to which our list of ion channels with the signature of positive selection are associated, we performed an analysis using the mammalian phenotype ontology database (Blake et al. 2003; Eppig et al. 2015) as implemented in the Enrichr platform (https://maayanlab.cloud/Enrichr/) (Xie et al. 2021).

In this analysis, we compared our gene list of positively selected ion channels (query list) against the gene list of the platform (subject list), which has 9767 genes grouped in 4601 phenotypical categories. The platform calculates the adjusted p-values from all raw p-values using the FDR procedure, and we considered only the phenotype categories with adjusted p-values equal to or below 0.01.

### Structural methods

Protein structure homology modeling was performed using the SWISS-MODEL server (https://swissmodel.expasy.org/) (Waterhouse et al. 2018). Structural figures were prepared using PyMOL Molecular Graphics System, Version 2.0.6 Schrödinger, LLC. Ligand-protein interaction diagrams were performed with LigPlot^+^ v.2.2 (Laskowski and Swindells 2011).

## Results and Discussion

### Ion channel annotation and homology inference

In this work, we studied the evolution of ion channels in cetaceans. To do this, we designed a bioinformatic pipeline (Fig. 1) to annotate the ion channel repertoire from the proteome of different mammalian species. Based on our analyses, we found that cetaceans possess fewer protein-coding genes than non-cetacean mammals (18845.5 ± 977.29 vs. 21396.22 ± 1218.75, unpaired one-tailed t-test with d.f.=14.88; t-statistic = - 4.94 and p-value = 9.1e-5) (Fig. 2). This result is consistent with other studies in which a reduction in gene copy number in cetaceans and other groups is associated with evolutionary innovations (Feng et al. 2014; Nery et al. 2014; Sun et al. 2017; Huelsmann et al. 2019; Helsen et al. 2020; McGowen et al. 2020; Cabrera et al. 2021; Randall et al. 2022; Zheng et al. 2022; Osipova et al. 2023; Pinto et al. 2023). We annotated, on average, 192.56 ion channels in the genomes of the species included in our taxonomic sampling, representing 0.96% of the protein-coding genes (Fig. 2). The smallest number of ion channels (177) was obtained for the bottlenose dolphin (*Tursiops truncatus*), while the largest number (226) was in humans (*Homo sapiens*) (Fig. 2). Our results are comparable, to what is reported in the literature: 235 ion channels for humans (*Homo sapiens*) and 231 for the mouse (*Mus musculus*) (Jegla et al. 2009). According to our results, on average, cetaceans possess a higher proportion of annotated ion channels in their genomes than the non-cetacean mammals (9.95 ± 0.38 vs. 0.92 ± 0.61, unpaired one-tailed t-test with d.f.=13.587; t-statistic = 2.933 and p-value = 0.005) (Fig. 2). Although the literature contains abundant examples of gene loss reported for cetaceans (see references above), there are also examples in which cetaceans expanded their gene repertoire. For instance, Holthaus et al. (2021) report that a subtype of small proline-rich proteins has expanded in cetaceans (Holthaus et al. 2021). Genes related to tumor suppression, cell cycle checkpoint, cell signaling, and proliferation have also expanded their repertoire in cetaceans (Tollis et al. 2019; Tejada-Martinez et al. 2021).

**Figure 2.**
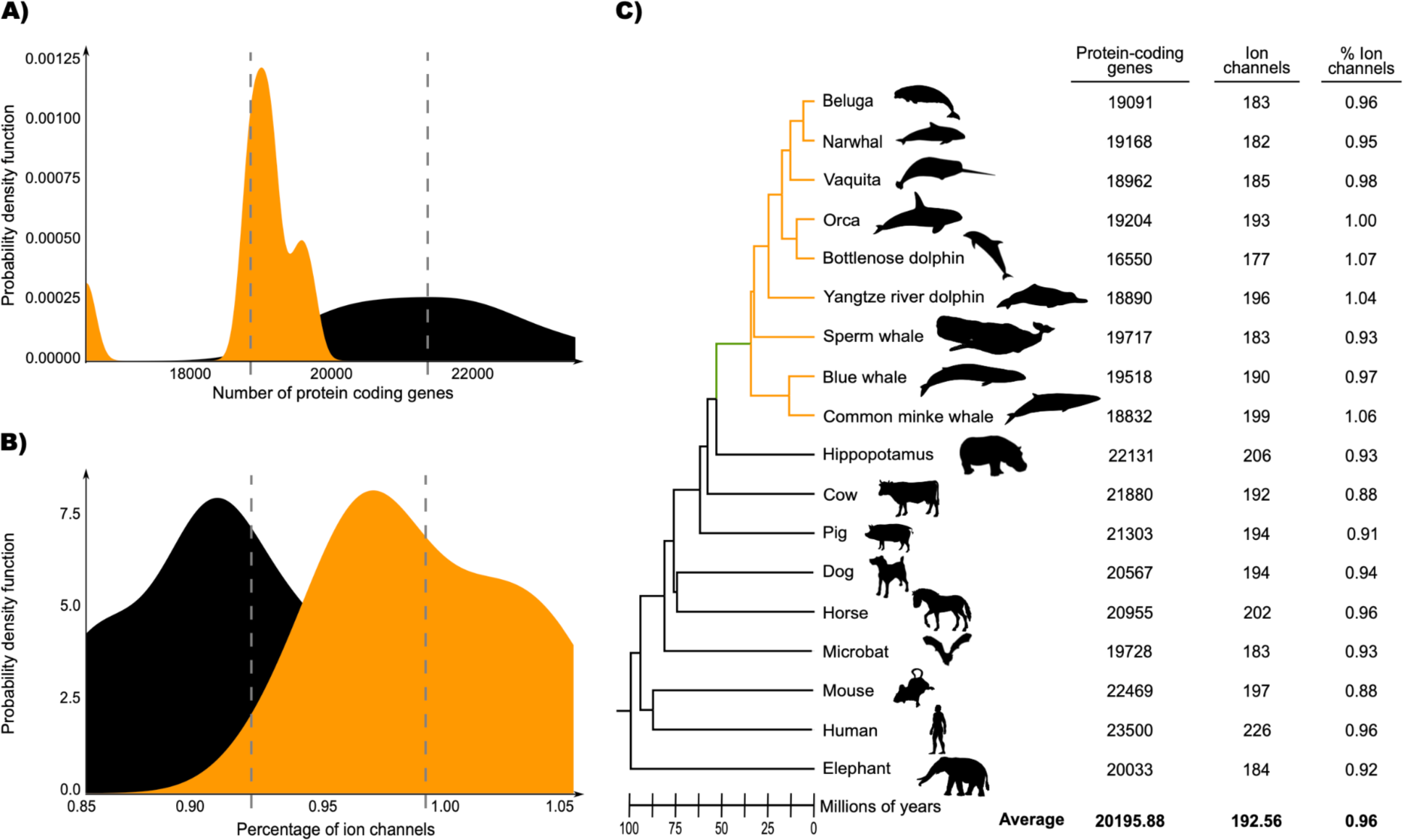
Comparison between averages of cetaceans (orange) versus non-cetaceans (black) in terms of the number of protein-coding genes (A) and percentage of ion channels among them (B). The Probability Density Functions were inferred through Kernel Density Estimation from the data processed from the genome annotations of the species included in our sampling. C) phylogenetic tree showing the species included in addition to the number of protein-coding genes, ion channels, and the percentage of ion channels. The green branch denotes the stem cetacea, while the orange branches are the crown cetacea. The divergence times were obtained from the Timetree 5 database (Kumar et al. 2022). Phylogenetic relationships were obtained from the literature (McGowen et al. 2019; Upham et al. 2019). Silhouette images were downloaded from PhyloPic (CC0 1.0 Universal Public Domain Dedication, https://creativecommons.org/publicdomain/zero/1.0/, http://phylopic.org/).

Through the homology inference procedure, we obtained 112 orthologous groups of putative ion channels present in all cetacean species. Additionally, 209 hierarchical orthologous groups containing two or more sequences from different species were inferred.

### Ion channels with the signature of positive selection are related to heart, locomotion, vision and neurological physiology

Using site analyses as implemented in PAML, we tested 112 orthologous groups of ion channels in which all cetacean species were present. According to our results, we retrieved 39 orthologous groups that were significant in both comparisons (M1vs M2 and M7 vs M8). After performing an enrichment analysis of the genes with the signal of positive selection against the mammalian phenotype ontology database (Eppig et al. 2015), we recovered categories related to phenotypes linked to the group of genes that are well known to have been modified due to the aquatic transition (e.g., circulatory, locomotor, and visual systems, among others). Comparable results have been obtained in previous studies where groups of genes related to specific phenotype are studied (Tollis et al. 2019; Tejada-Martinez et al. 2021; Silva et al. 2023), also in genome-wide studies (McGowen et al. 2012; Nery, González, et al. 2013; Sun et al. 2013; Park et al. 2015).

Our study found a category related to heart physiology. We identified categories such as “decreased cardiac muscle contractility”, “irregular heartbeat”, and “atrial fibrillation”, which align with previous literature on changes in heart function during diving (Fig. 3 and Supplementary Table S2) (Scholander 1940; Irving et al. 1941; Williams et al. 2015; Goldbogen et al. 2019). One of the main challenges for aquatic living is coping with the acute hypoxia that occurs during extended breath-holding. To overcome the physiological effects of deep dives, remarkable adaptations have evolved, including significant decreases in heart rate and redistribution of blood flow to essential aerobic tissues (Kooyman and Ponganis 1998). Although the basic structure of the cetacean heart is similar to that of other mammals, based on our findings we hypothesized the importance of ion channels, among other proteins, in adapting to diving. Thus, besides the well-documented morphological changes in the cetacean heart (Tarpley et al. 1997; Latorre et al. 2022) mostly attributed to diving adaptations, our results show adaptive changes in genes encoding ion channels, which are fundamental for their physiological divergence.

**Figure 3.**
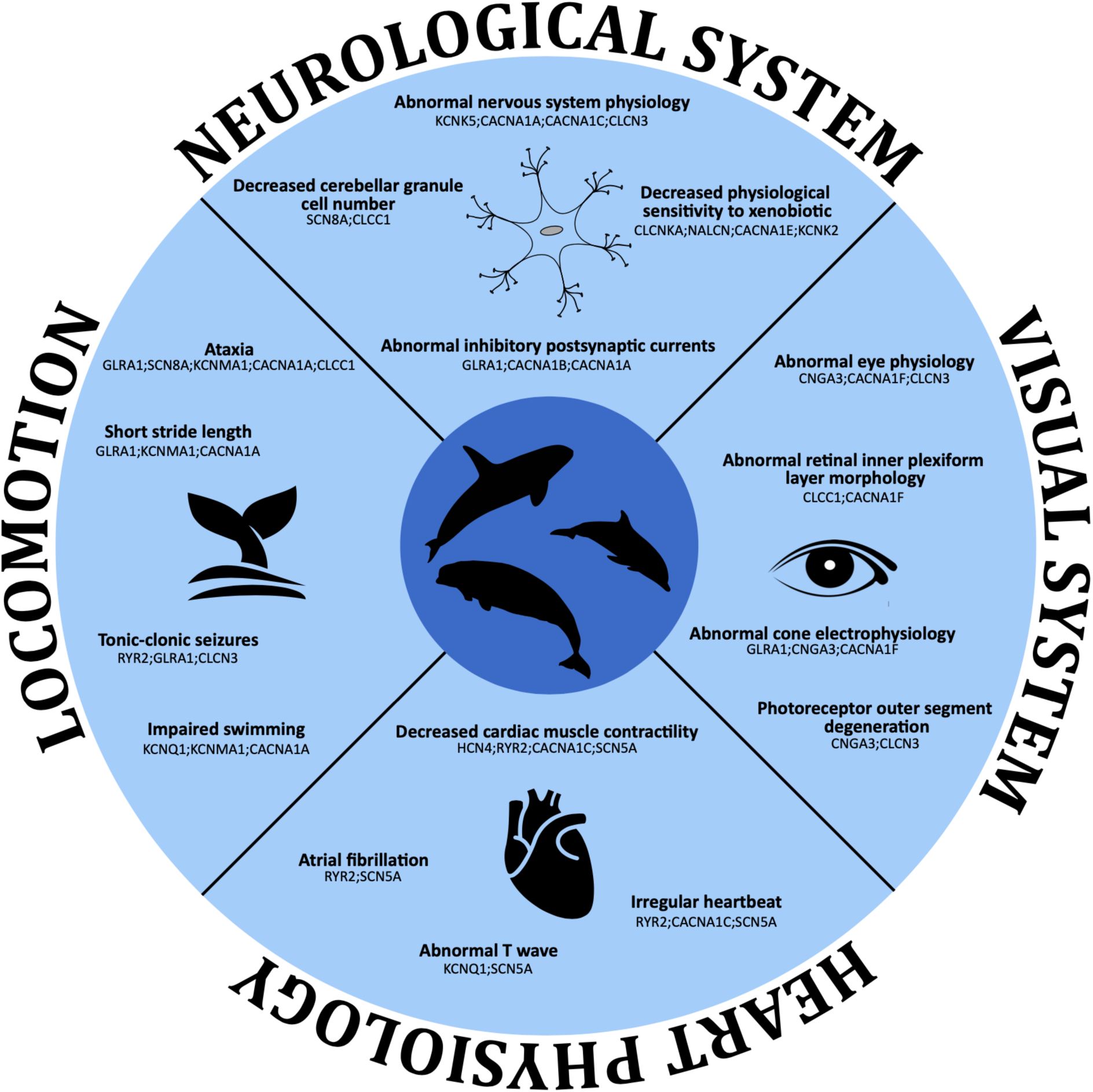
Enriched categories related to heart physiology, locomotion, visual system and neurological system in cetaceans based on the MGI mammalian phenotype ontology database using the Enrichr platform (https://maayanlab.cloud/Enrichr/). For further details see Supplementary Tables S1, S2, S3, S4 and S5.

Among the genes associated with the recovered categories related to heart physiology (Fig. 3 and Supplementary Table S2), the Sodium Voltage-Gated Channel Alpha Subunit 5 (SCN5A) gene is the most frequent. The SCN5A gene encodes the alpha subunit of the sodium channel Na_V_1.5, mainly expressed in the heart (Uhlen et al. 2010). This protein mediates the voltage-dependent sodium ion permeability of the cardiac muscle, playing an essential role in regulating cardiac electrophysiological function (Li et al. 2018). In addition, mutations in this ion channel cause a broad range of electrical disorders and structural abnormalities in humans (Li et al. 2018; D. Jiang et al. 2020; Rivaud et al. 2020). Past studies have also found this ion channel under positive selection in cetaceans (Sun et al. 2013).

Although the Na_V_1.5 and its paralog, the Sodium Voltage-Gated Channel Alpha Subunit 9 (Na_V_1.7) sodium channel have similar selectivity filters (Shen et al. 2019; D. Jiang et al. 2020), the Na_V_1.5 channel has an affinity about two orders of magnitude lower for tetrodotoxin (TTX) (Sunami et al. 2000; Walker et al. 2012), a sodium channel blocker that is found in a variety of marine animals (e.g., pufferfish, horseshoe crabs, blue-ringed octopus, gastropods, starfish, among others) (Jal and Khora 2015). This difference is due to a single amino acid substitution, where the Na_V_1.5 channel has a cysteine at position 373 instead of a tyrosine (Fig. 4). As expected, all Na_V_1.7 channels of cetaceans possess a tyrosine amino acid at a structurally related position, Y362 in human Na_V_1.7 (Fig. 4), making it susceptible to TTX blockage. Interestingly, besides the bottlenose dolphins, the Na_V_1.5 channel of the toothed whales (Odontoceti) species included in our study possesses a tyrosine instead of cysteine (Fig. 4), making it potentially sensitive to TTX. Furthermore, three-dimensional protein structure modeling of sperm whale Na_V_1.5 and Na_V_1.7 shows a conserved spatial arrangement of amino acid residues that are important for TTX binding on human Na_V_1.7, including Y362 that establishes a ϰ-cation interaction with the guanidinium group of TTX, an interaction that is not possible by C373 of human Na_V_1.5 (Fig. 5). To strengthen our conclusions, we retrieved additional Na_V_1.5 sequences from toothed whales to see if they also have tyrosine instead of cysteine. In our new searches, we recovered the Na_V_1.5 ion channel of the long-finned pilot whale (*Globicephala melas*), Pacific white-sided dolphin (*Lagenorhynchus obliquidens*) and Yangtze finless porpoise (*Neophocaena asiaeorientalis*). We found a tyrosine residue in all cases, providing further support to our results (Fig. 4). We also found a different species of the genus *Tursiops*, the Indo-Pacific bottlenose dolphin (*Tursiops aduncus*), and following the trend we are describing, it has a cysteine residue (Fig. 4). Finally, to be sure that this cysteine to tyrosine replacement only occurred in toothed whales, we also retrieved additional Na_V_1.5 sequences from baleen whales (Mysticeti). We added two more species, the humpback whale (*Megaptera novaeangliae*) and the gray whale (*Eschrichtius robustus*). Both species have a cysteine residue, confirming our initial conclusion (Fig. 4). Thus, the amino acid substitution from cysteine to tyrosine occurred in the last common ancestor of toothed whales (Odontoceti), which lived between 34 and 33 million years ago (Kumar et al. 2022). However, in the last common ancestor of the genus *Tursiops*, it was reversed (Fig. 4). This is interesting because documentarists have been able to record dolphins “playing” with pufferfish, with the threat of being exposed to TTX that the pufferfish releases when it feels threatened. However, since the toxin is released into the water, it seems to be not in lethal doses, and it has been proposed it could cause a trance-like state. This could be possible because pufferfish mainly accumulate TTX in the skin (Zhang et al. 2020), especially when they are young (Tatsuno et al. 2013), an organ in direct contact with the mouth of the dolphin. In fact, it has been observed that dolphins behave differently after “playing” with the pufferfish. Therefore, this behavior could be related to the reversal in the amino acid substitution, which makes it plausible that the Na_V_1.5 ion channel of the species of the genus *Tursiops* has a lower affinity for TTX, like all mammals. Thus, besides having mutations associated with human channelopathies, the Na_V_1.5 channel of a subset of cetaceans, toothed whales (Odontoceti), is predicted to have a higher affinity to TTX than other mammals. Further lab experiments are the next logical step to reveal the functional divergence of the Na_V_1.5 in this particular group of mammals.

**Figure 4.**
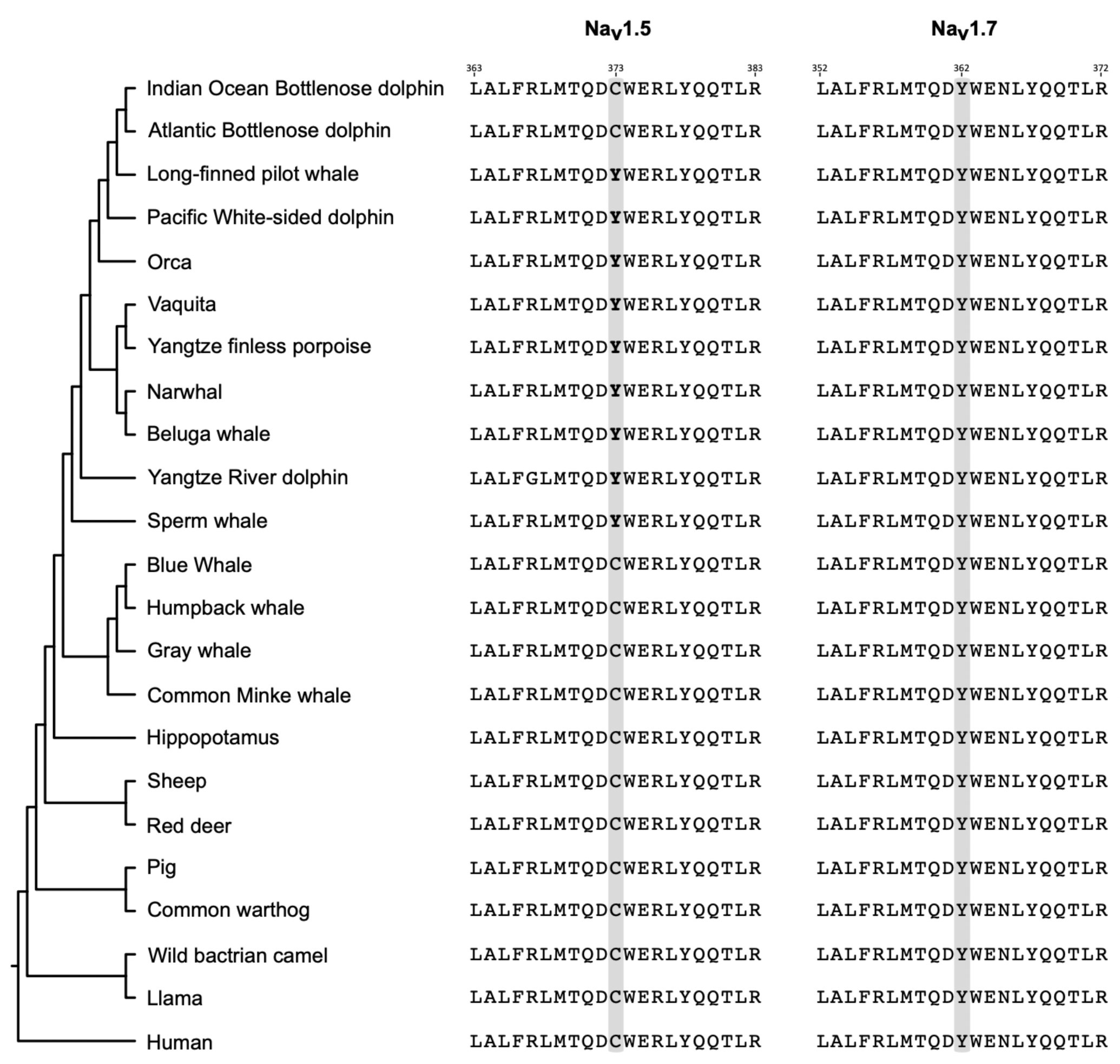
Amino acid alignment of a segment between the transmembrane segments 5 and 6 of the NaV1.5 and NaV1.7 ion channels containing the amino acid positions responsible for the sensitivity to TTX (shaded in gray). The numbering of the amino acids is with respect to the human sequences (NaV1.7, NM_001365536.1; NaV1.5, NM_001099404.2). Phylogenetic relationships were obtained from the literature (McGowen et al. 2019; Upham et al. 2019). In bold are the amino acids tyrosine (Y) in the NaV1.5 ion channels of toothed whales (Odontoceti), which would make them more sensitive to TTX.

**Figure 5.**
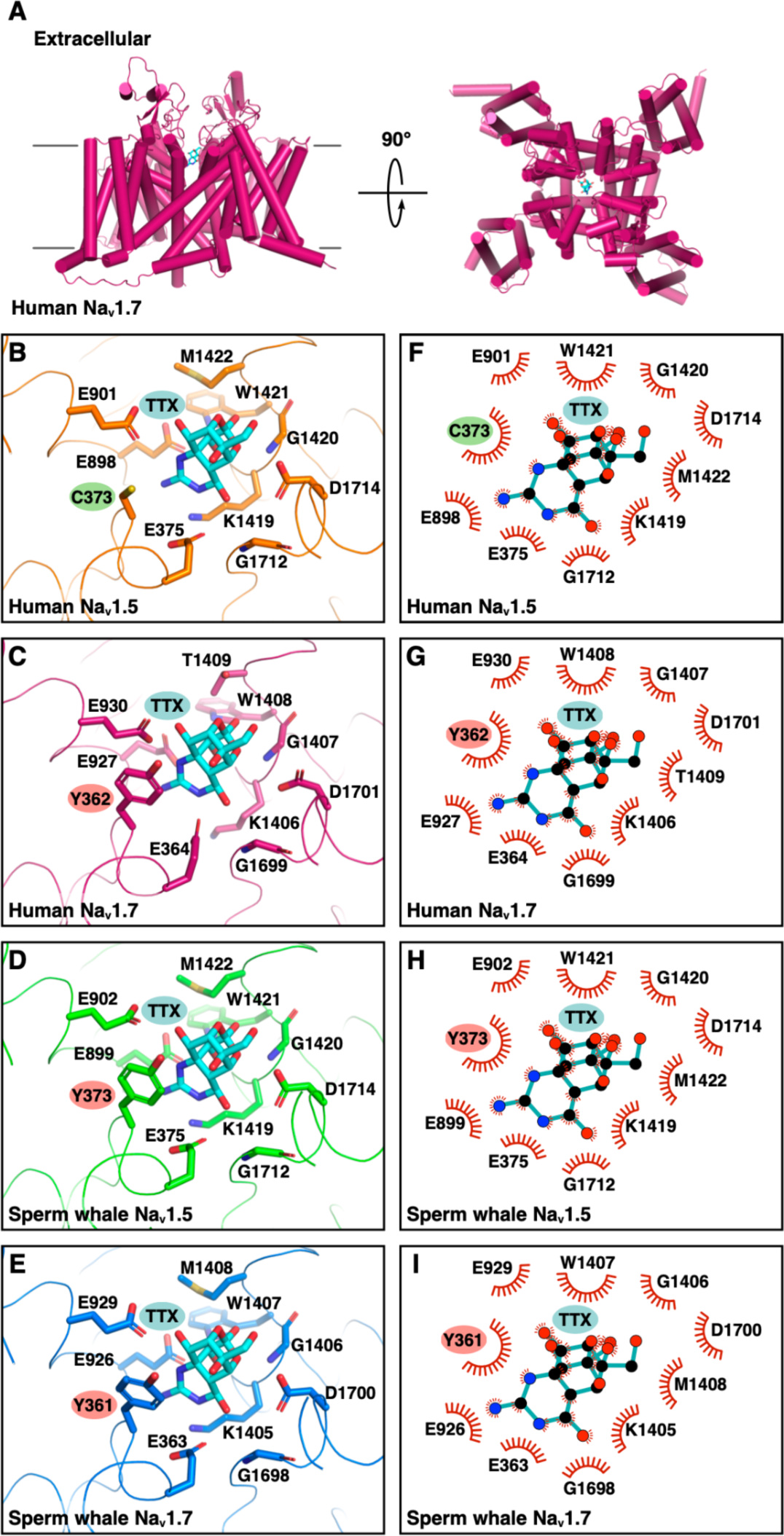
Comparison of the binding site for TTX in human and sperm whale NaV1.5 and NaV1.7. A) Cartoon representation (alpha-helices shown as cylinders) of human NaV1.7 colored magenta, and stick model of TTX with carbon atoms colored cyan (PDB code 6J8I showing only chain B and TTX). B-E) Ribbon representation of the TTX binding site on the indicated sodium channels, and stick representation of sodium channel TTX binding site amino acid residues with carbon atoms colored orange (B; human NaV1.5; PDB code 6LQA), magenta (C; human NaV1.7; PDB code 6J8I), green (D; sperm whale NaV1.5; model generated by SWISS-MODEL), or blue (E; sperm whale NaV1.7; model generated by SWISS-MODEL), and of TTX with carbon atoms colored cyan. Highlighted are the positions of C373 in human NaV1.5, of Y362 in human NaV1.7 which is critical for binding via a ϰ-cation interaction with the 1,2,3-guanidinium group of TTX, and of Y373 and Y361 of sperm whale NaV1.5 and NaV1.7, respectively, predicted to interact with TTX via a similar p-cation interaction. The carbonyl and amino groups of glycine residues are depicted for better visualization. F-I) Two-dimensional, schematic representation of the position of TTX shown in B-E using LigPlot^+^.

The Ryanodine Receptor 2 (RYR2) gene, the second most frequent gene in Table 1, encodes for a protein that is one of the components of the largest ion channel known to date. It is mainly expressed in the heart (Uhlen et al. 2010), and its primary function is controlling Ca^+2^ release from the sarcoplasmic reticulum throughout the cardiac cycle (Fowler and Zissimopoulos 2022). Mutations in this channel have been associated with Arrhythmogenic Right Ventricular Dysplasia, familial, 2 (ARVD2), Ventricular tachycardia, catecholaminergic polymorphic, 1 (CPVT1), Ventricular arrhythmias due to cardiac ryanodine receptor calcium release deficiency syndrome (VACRDS), among others (Sleiman et al. 2021).

We also retrieved categories related to locomotion (Fig. 3 and Supplementary Table S3), one of the phenotypes significantly modified in cetaceans, with profound changes in the body plan due to the aquatic transition. Unlike terrestrial mammals, cetaceans have elongated bodies and absent hindlimbs. The forelimb was modified into a flipper, and vertical movements of the tail accomplished locomotion. Moreover, moving into the water column imposes a highly energetic cost requiring a greater force than moving into the terrestrial environment, which translates into a need for stronger muscle contraction (Berta et al. 2005). In stronger muscle contraction, ion channels play a fundamental role by initiating the influx of ions, including sodium transport, into the cells. This result agrees with Sun et al. (2013), which reported the signature of positive selection in genes related to motor activity and muscle contraction in cetaceans. The two most frequently mentioned ion channels are the Calcium Voltage-Gated Channel Subunit Alpha1 A (CACNA1A) and Glycine Receptor Alpha 1 (GLRA1). Thus, like the heart case previously described, given that marine mammal muscle architecture is similar to most other mammals (Würsig et al. 2017), the fact that we recovered several ion channels related to locomotion highlights their fundamental role in physiological divergence.

The senses of cetaceans have radically changed in association with aquatic living (De Vreese et al. 2023). The cetacean visual system has been modified to meet the challenges of the aquatic way of life, in which ion channels seem to be an important factor (Fig. 3 and Supplementary Table S4). Unlike many terrestrial mammals, cetaceans have eyes positioned laterally on their heads, which provide a panoramic view of their surroundings. This lateral placement is crucial for wide-range detection of both predators and preys in their vast aquatic habitat (Mass and Supin 2018). Although cetacean eyesight is generally considered less acute than that of terrestrial mammals, it includes specific adaptations for underwater vision. For example, a key adaptation is the higher density of rod cells in their retinas, implying enhanced vision in low-light conditions prevalent in deeper waters (Peichl et al. 2001). Also, cetaceans lack a fovea, a slight depression within the retina, a region associated with visual acuity in other mammals (Dawson et al. 1987). This absence suggests an evolutionary trade-off, where the demands of aquatic vision have shaped a different visual acuity strategy. Furthemore, the evolutionary history of cetacean opsin genes, responsible for light detection, reflects an adaptation to the marine light environment (Fasick and Robinson 2016; Mass and Supin 2018).

We also identified a category related to the neurological system (Fig. 3). This category encompasses all aspects of phenotypic changes within the nervous system attributed to the diverse sensory adaptations now thoroughly documented as originating from the transition to aquatic life (Ridgway 1988; Eldridge et al. 2022; Racicot 2022; De Vreese et al. 2023). These phenotypic changes include modifications in brain morphology, sensory processing, and motor control to support their aquatic lifestyle (Thewissen 2018). Accordingly, we retrieved categories associated with abnormal nervous system physiology, brain morphology, and excitatory postsynaptic potential, among others (Fig. 3 and Supplementary Table S5). The most frequent genes in this category are CACNA1A and SCN8A. Associated with this category, we also retrieved ion channels related to hearing, although not among the most significantly enriched categories. We identified ion channels associated with ear and otolith morphology anomalies, among other findings. Accompanying the specialized hearing capabilities in this mammalian group, neuroanatomical alterations affecting the inner ear and cranial nerves have been documented (De Vreese et al. 2023).

### Accelerated gene turnover rate in cetaceans

The availability of whole-genome sequences has revealed that variation in gene copy number is abundant in nature (Schrider and Hahn 2010) and related to the origin of phenotypic diversity. For example, a survey of more than 9,000 gene families in primates suggested that humans possess faster gene turnover than other mammals (Hahn et al. 2007). In this study, the authors found several expansions (e.g., centaurin gamma gene family) in the lineage leading to humans that are related to the unique attributes of our species (e.g., enlarged brains) (Hahn et al. 2007), establishing a link between copy number variation and evolutionary innovations.

In our results, we also found variation in the gene turnover rate (Fig. 6). Our models estimating different λ parameters for cetaceans, as a total group and non-cetacean mammals, showed that the first group possesses a rate of evolution (λ_C_ = 0.0023) 2.87 times faster compared to non-cetacean mammals (λ_o_ = 0.0008; Fig. 6). In the second model the estimated λ parameter for the crown group cetacea (λ_C_ = 0.0025) was 3.12 times faster in comparison to non-cetacean mammals (λ_0_ = 0.0008) and 12.5 times faster than the value estimated for the last common ancestor of cetaceans (λ_anc_ = 0.0002) (Fig. 6), suggesting that most of the variation in gene copy number occurred during the radiation of the group. Similar results were obtained when tumor suppressor genes were analyzed (Tejada-Martinez et al. 2021).

**Figure 6.**
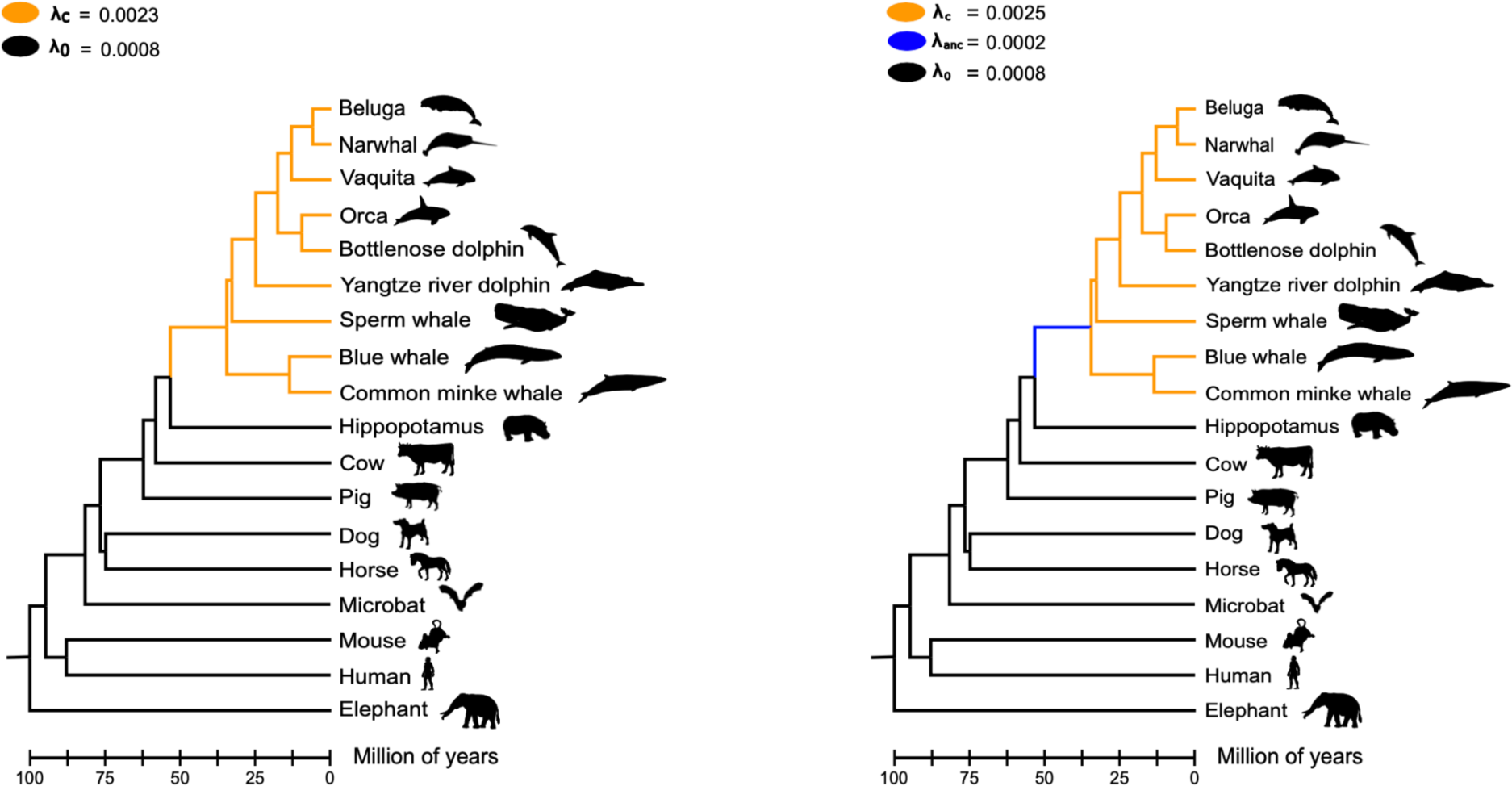
Gene turnover rates of ion channels. The first model (left panel) estimated the rate of evolution (λ) of ion channels for cetaceans as a total group (orange branches) and for non-cetacean mammals (black branches). Under this model, the λ value for cetaceans is more than two times faster than non-cetacean mammals. The second model (right panel) estimated λ values for the last common ancestor of cetaceans (blue branch), for the crown group cetacea (orange branches), and for non-cetacean mammals (black branches).

According to our estimates, the number of copies of ion channel genes in cetaceans varies between zero and 18. We found four hierarchical orthologous groups in which all cetacean species have no copies (ASIC5, CLDND1, KCNMB3, and PKD1L1). In these cases, we double-checked the information and found different situations. In the case of the acid-sensing ion channel 5 (ASIC5), we found it in the FASTA file containing all protein-coding genes of baleen whales (Mysticeti), but it was predicted to have less than two transmembrane segments, so it was not included in the next step in our pipeline. A similar situation occurred for the calcium-activated potassium channel subunit beta-3 (KCNMB3) and Claudin domain containing 1 (CLDND1) genes. The polycystic kidney disease protein 1-like 1 (PKD1L1) gene was only found in Orca, but it was not predicted to have any transmembrane segment. We also noticed that the PKD1L1 sequence associated with the accession number (XP_033267232.1) was removed from NCBI. According to ENSEMBL, no cetacean species was predicted to have an ortholog of the human PKD1L1 gene. Further, a comparison of the genomic region of the human (*Homo sapiens*), which possesses the PKD1L1 gene, with the corresponding chromosomal region in the sperm whale (*Physeter macrocephalus*), common minke whale (*Balaenoptera acutorostrata*), and vaquita (*Phocoena sinus*) suggests that the PKD1L1 gene is not present in the cetacean lineage (Fig. 7). It is worth noting that traces of the PKD1L1 gene are present in the sperm whale and vaquita genomes (Fig. 7). In figure 7 we also included the comparison with the horse (*Equus caballus)*, a species that share a common ancestor with humans at the same age as cetaceans, to show the conservation pattern of the chromosomal region harboring the PKD1L1 gene (Fig. 7). So, it is highly probable that this gene is not present in the cetacean genome. This result agrees with the study of Turakhia et al. (2020); however, they also show that gene loss is not a cetacean-specific evolutionary event, as they did not find the PKD1L1 gene in other cetartiodactyla species (e.g., alpaca, Bactrian camel, goat, sheep, Tibetan antelope, and cow) (Turakhia et al. 2020). In agreement, the cow and pig genome lack the PKD1L1 hierarchical orthologous group. Thus, this result suggests that the deletion of the PKD1L1 gene occurred in the last common ancestor of cetartiodactyla. PKD1L1 is a member of the TRP gene family, mainly expressed in the testis and heart (Yuasa et al. 2002; Cabezas-Bratesco et al. 2022). It also has functions related to establishing left-right asymmetry in positioning and patterning internal organs and associated vasculature by forming heteromeric channels with PKD2, functioning as sensors of the nodal flow (Pennekamp et al. 2002; Field et al. 2011; Norris 2012; Grimes et al. 2016; Esarte Palomero et al. 2023). However, this gene loss does not translate into functional consequences in cetaceans and even-toed ungulates, probably due to a certain degree of redundancy that could serve as a backup with functionally overlapping family members (Nowak et al. 1997; Félix and Barkoulas 2015; Albalat and Cañestro 2016).

**Figure 7.**
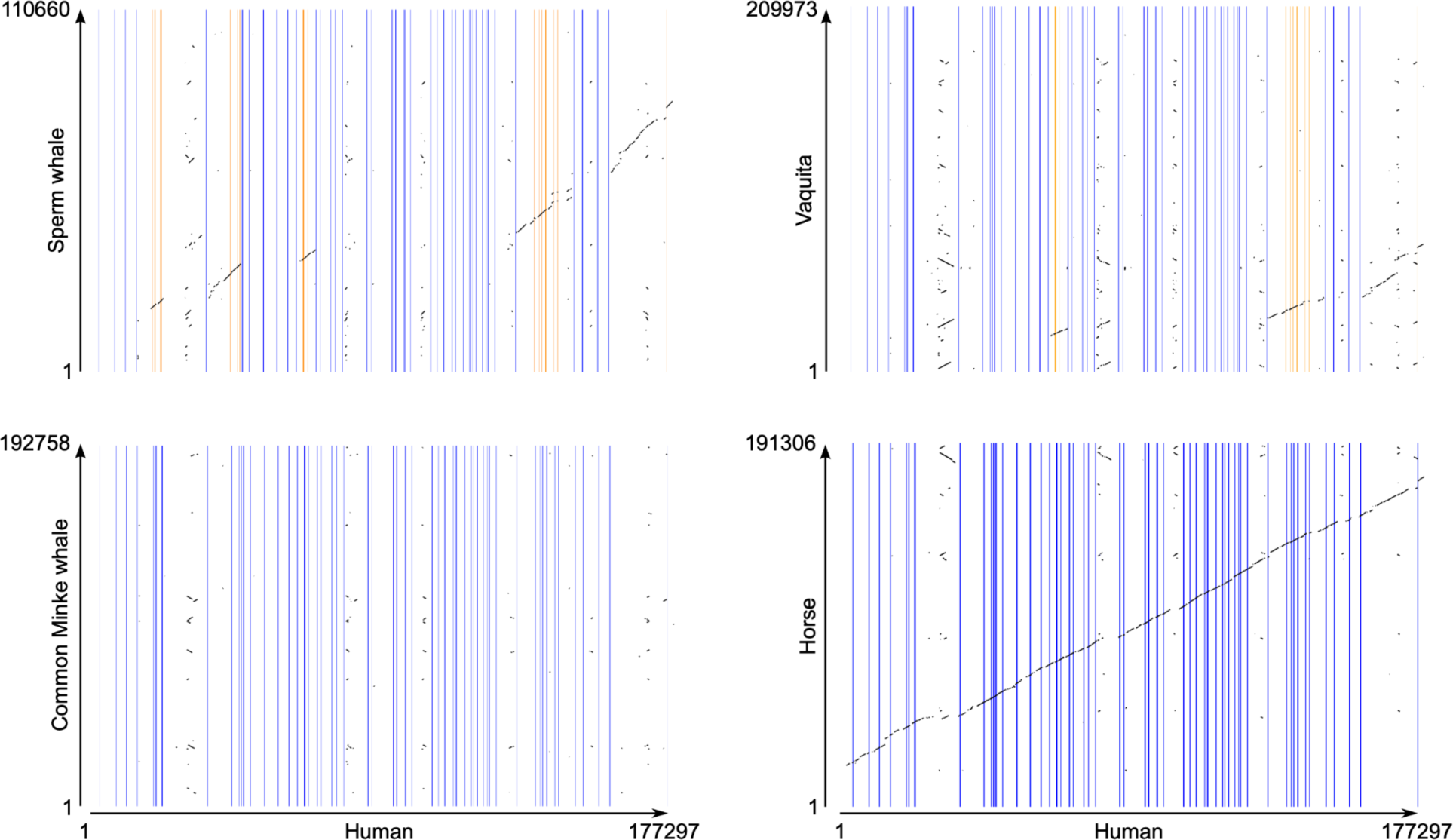
Pairwise dot-plot comparison of the genomic region containing the polycystic kidney disease protein 1-like 1 (PKD1L1) gene of the human (*Homo sapiens*) with the corresponding region in the sperm whale (*Physeter macrocephalus*), common minke whale (*Balaenoptera acutorostrata*), vaquita (*Phocoena sinus*), and horse (*Equus caballus*). Vertical lines denote exons, and regions in between are introns. Orange vertical lines indicate exons that are still present in the species that lost the PKD1L1 gene.

The hierarchical orthologous group that contained the highest number of copies corresponded to the transmembrane protein 37 (TMEM37), a gamma subunit of voltage-gated calcium channels, with 18 copies. In the case of the non-cetacean mammals in our sampling, this hierarchical orthologous group has nine copies, suggesting that the 18 copies represent a cetacean-specific expansion. After that hierarchical orthologous group with 18 gene copies, there is a group with 12 copies (P2RX4), another with 11 copies (KCNK13) and two groups with 10 copies (CLCN4 and KCNH4). In all cases, the non-cetacean mammals in our sampling have fewer copies than cetaceans, six for P2RX4 and nine for KCNK13, CLCN4 and KCNH4.

A more detailed assessment of the hierarchical orthologous groups provides a better panorama of the evolutionary trend in which cetacean species possess fewer ion channels than the non-cetacean mammals in our sampling. Our analysis found that out of the 209 hierarchical orthologous groups inferred, cetaceans possess more gene copies for 51 groups. On the other hand, in 94 hierarchical orthologous groups, both groups have the same number of gene copies, and, in 64 hierarchical orthologous groups, non-cetacean mammals possess more copies than cetaceans.

## Conclusions

In this work, we designed a bioinformatic pipeline that identifies the entire repertoire of ion channels of any species. In our case, we used it to study the evolution of ion channels in cetaceans, a mammalian group that, due to the conquest of the aquatic environment, has extensively modified physiological axes in which ion channels play a significant role. Our results indicate that cetaceans have on average, fewer protein-coding genes and a higher percentage of annotated ion channels than non-cetacean mammals. Furthermore, most of the genes with the signal of positive selection are related to heart, locomotion, visual and neurological phenotypes, consistent with previous studies. The Na_V_1.5 channel of mammals is about two orders of magnitude less sensitive to TTX than NaV1.7. This difference is due to a cysteine residue instead of a tyrosine in a specific position. However, our work shows that most species of toothed whales (Odontoceti) possess a tyrosine amino acid residue in that particular position of the Na_V_1.5 channel, making them potentially sensitive to TTX, similar to Na_V_1.7. However, it is important to recall that the effect of an amino acid replacement depends on the genetic background. So, future studies should be directed to evaluate the biochemical/biophysical performance of the Na_V_1.5 channel of toothed whales, especially their sensitivity to TTX. Finally, the natural experiment that cetaceans represent in the Tree of Life provides an excellent model to advance our understanding of the genetic bases of phenotypic diversity. The number of genomes that exist today and those that are yet to come will make the field of evolutionary genomics a significant contributor to different disciplines in biology and medicine.

## Funding

We thank two reviewers for their constructive criticism and Gavin Douglas for such great work as a PCI genomics recommender. This work was supported by the Fondo Nacional de Desarrollo Científico y Tecnológico from Chile, FONDECYT 1210471 to JCO, 1231357 to GR and 1211481 to GAM. FONDEQUIP grant EQM160063 to GR.

The Millennium Nucleus of Ion Channel-Associated Diseases is a Millennium Nucleus of the Millennium Scientific Initiative, National Agency of Research and Development (ANID, NCN19_168), Ministry of Science, Technology, Knowledge, and Innovation, Chile to JCO and GR. This study was also supported by Coordination for the Improvement of Higher Education Personnel—Brasil (CAPES)—Finance Code 001 and FAPESP (2015/18269-1) to MFN.

## Conflict of Interest disclosure

The authors declare they have no conflict of interest relating to the content of this article.

## Data, script, code, and supplementary information availability

Data and supplementary material are available online at https://github.com/opazolab/Cetacean_ion_channels

## Notes

https://github.com/opazolab/Cetacean_ion_channels

### Competing Interest Statement

The authors have declared no competing interest.

### Summary of Updates

Just a little change to the title

## References

Ackerman MJ. 2004. Cardiac channelopathies: it’s in the genes. Nat. Med. 10:463–464.

Ackerman MJ, Clapham DE. 1997. Ion channels--basic science and clinical disease. N. Engl. J. Med. 336:1575–1586.

Albalat R, Cañestro C. 2016. Evolution by gene loss. Nat. Rev. Genet. 17:379–391.

Altenhoff AM, Train C-M, Gilbert KJ, Mediratta I, Mendes de Farias T, Moi D, Nevers Y, Radoykova H-S, Rossier V, Warwick Vesztrocy A, et al. 2021. OMA orthology in 2021: website overhaul, conserved isoforms, ancestral gene order and more. Nucleic Acids Res. 49:D373–D379.

Benjamini Y, Hochberg Y. 1995. Controlling the false discovery rate: A practical and powerful approach to multiple testing. J. R. Stat. Soc. 57:289–300.

Berta A, Sumich JL, Kovacs KM. 2005. Marine Mammals: Evolutionary Biology. Elsevier

Blake JA, Richardson JE, Bult CJ, Kadin JA, Eppig JT, Mouse Genome Database Group. 2003. MGD: the Mouse Genome Database. Nucleic Acids Res. 31:193–195.

Bondareva O, Petrova T, Bodrov S, Gavrilo M, Smorkatcheva A, Abramson N. 2023. How voles adapt to subterranean lifestyle: Insights from RNA-seq. Front. Ecol. Evol. [Internet] 11. Available from: https://www.frontiersin.org/articles/10.3389/fevo.2023.1085993/full

Cabezas-Bratesco D, Mcgee FA, Colenso CK, Zavala K, Granata D, Carnevale V, Opazo JC, Brauchi SE. 2022. Sequence and structural conservation reveal fingerprint residues in TRP channels. Elife [Internet] 11. Available from: 10.7554/eLife.73645

Cabrera AA, Bérubé M, Lopes XM, Louis M, Oosting T, Rey-Iglesia A, Rivera-León VE, Székely D, Lorenzen ED, Palsbøll PJ. 2021. A genetic perspective on cetacean evolution. Annu. Rev. Ecol. Evol. Syst. 52:131–151.

De Vreese S, Orekhova K, Morell M, Gerussi T, Graïc J-M. 2023. Neuroanatomy of the Cetacean Sensory Systems. Animals (Basel) [Internet] 14. Available from: 10.3390/ani14010066

Eldridge SA, Mortazavi F, Rice FL, Ketten DR, Wiley DN, Lyman E, Reidenberg JS, Hanke FD, DeVreese S, Strobel SM, et al. 2022. Specializations of somatosensory innervation in the skin of humpback whales (Megaptera novaeangliae). Anat. Rec. 305:514–534.

Eppig JT, Blake JA, Bult CJ, Kadin JA, Richardson JE, Mouse Genome Database Group. 2015. The Mouse Genome Database (MGD): facilitating mouse as a model for human biology and disease. Nucleic Acids Res. 43:D726–D736.

Esarte Palomero O, Larmore M, DeCaen PG. 2023. Polycystin Channel Complexes. Annu. Rev. Physiol. 85:425–448.

Fasick JI, Robinson PR. 2016. Adaptations of cetacean retinal pigments to aquatic environments. Front. Ecol. Evol. [Internet] 4. Available from: http://journal.frontiersin.org/Article/10.3389/fevo.2016.00070/abstract

Félix M-A, Barkoulas M. 2015. Pervasive robustness in biological systems. Nat. Rev. Genet. 16:483–496.

Feng P, Zheng J, Rossiter SJ, Wang D, Zhao H. 2014. Massive losses of taste receptor genes in toothed and baleen whales. Genome Biol. Evol. 6:1254–1265.

Field S, Riley K-L, Grimes DT, Hilton H, Simon M, Powles-Glover N, Siggers P, Bogani D, Greenfield A, Norris DP. 2011. Pkd1l1 establishes left-right asymmetry and physically interacts with Pkd2. Development 138:1131–1142.

Fowler ED, Zissimopoulos S. 2022. Molecular, Subcellular, and Arrhythmogenic Mechanisms in Genetic RyR2 Disease. Biomolecules [Internet] 12. Available from: 10.3390/biom12081030

Giroud S, Chery I, Arrivé M, Prost M, Zumsteg J, Heintz D, Evans AL, Gauquelin-Koch G, Arnemo JM, Swenson JE, et al. 2021. Hibernating brown bears are protected against atherogenic dyslipidemia. Sci. Rep. 11:18723.

Goldbogen JA, Cade DE, Calambokidis J, Czapanskiy MF, Fahlbusch J, Friedlaender AS, Gough WT, Kahane-Rapport SR, Savoca MS, Ponganis KV, et al. 2019. Extreme bradycardia and tachycardia in the world’s largest animal. Proc. Natl. Acad. Sci. U. S. A. 116:25329–25332.

Grimes DT, Keynton JL, Buenavista MT, Jin X, Patel SH, Kyosuke S, Vibert J, Williams DJ, Hamada H, Hussain R, et al. 2016. Genetic Analysis Reveals a Hierarchy of Interactions between Polycystin-Encoding Genes and Genes Controlling Cilia Function during Left-Right Determination. PLoS Genet. 12:e1006070.

Hahn MW, Demuth JP, Han S-G. 2007. Accelerated rate of gene gain and loss in primates. Genetics 177:1941–1949.

Han MV, Thomas GWC, Lugo-Martinez J, Hahn MW. 2013. Estimating gene gain and loss rates in the presence of error in genome assembly and annotation using CAFE 3. Mol. Biol. Evol. 30:1987–1997.

Helsen J, Voordeckers K, Vanderwaeren L, Santermans T, Tsontaki M, Verstrepen KJ, Jelier R. 2020. Gene Loss Predictably Drives Evolutionary Adaptation. Mol. Biol. Evol. 37:2989–3002.

Holthaus KB, Lachner J, Ebner B, Tschachler E, Eckhart L. 2021. Gene duplications and gene loss in the epidermal differentiation complex during the evolutionary land-to-water transition of cetaceans. Sci. Rep. 11:12334.

Houssaye A, Fish FE. 2016. Functional (Secondary) Adaptation to an Aquatic Life in Vertebrates: An Introduction to the Symposium. Integr. Comp. Biol. 56:1266–1270.

Huelsmann M, Hecker N, Springer MS, Gatesy J, Sharma V, Hiller M. 2019. Genes lost during the transition from land to water in cetaceans highlight genomic changes associated with aquatic adaptations. Sci Adv 5:eaaw6671.

Irving L, Scholander PF, Grinnell SW. 1941. Significance of the heart rate to the diving ability of seals. Journal of Cellular and Comparative Physiology [Internet] 18:283–297. Available from: 10.1002/jcp.1030180302

Jal S, Khora SS. 2015. An overview on the origin and production of tetrodotoxin, a potent neurotoxin. J. Appl. Microbiol. 119:907–916.

Jegla TJ, Zmasek CM, Batalov S, Nayak SK. 2009. Evolution of the human ion channel set. Comb. Chem. High Throughput Screen. 12:2–23.

Jiang D, Shi H, Tonggu L, Gamal El-Din TM, Lenaeus MJ, Zhao Y, Yoshioka C, Zheng N, Catterall WA. 2020. Structure of the Cardiac Sodium Channel. Cell 180:122– 134.e10.

Jiang M, Shi L, Li X, Dong Q, Sun H, Du Y, Zhang Y, Shao T, Cheng H, Chen W, et al. 2020. Genome-wide adaptive evolution to underground stresses in subterranean mammals: Hypoxia adaption, immunity promotion, and sensory specialization. Ecol. Evol. 10:7377–7388.

Kashio M, Tominaga M. 2022. TRP channels in thermosensation. Curr. Opin. Neurobiol. 75:102591.

Katoh K, Standley DM. 2013. MAFFT multiple sequence alignment software version 7: improvements in performance and usability. Mol. Biol. Evol. 30:772–780.

Kelley JL, Brown AP, Therkildsen NO, Foote AD. 2016. The life aquatic: advances in marine vertebrate genomics. Nat. Rev. Genet. 17:523–534.

Kim J-B. 2014. Channelopathies. Korean J. Pediatr. 57:1–18.

Kooyman GL, Ponganis PJ. 1998. The physiological basis of diving to depth: birds and mammals. Annu. Rev. Physiol. 60:19–32.

Krogh A, Larsson B, von Heijne G, Sonnhammer EL. 2001. Predicting transmembrane protein topology with a hidden Markov model: application to complete genomes. J. Mol. Biol. 305:567–580.

Kumar S, Suleski M, Craig JM, Kasprowicz AE, Sanderford M, Li M, Stecher G, Hedges SB. 2022. TimeTree 5: An Expanded Resource for Species Divergence Times. Mol. Biol. Evol. [Internet] 39. Available from: 10.1093/molbev/msac174

Laskowski RA, Swindells MB. 2011. LigPlot+: multiple ligand-protein interaction diagrams for drug discovery. J. Chem. Inf. Model. 51:2778–2786.

Latorre R, Graïc J-M, Raverty SA, Soria F, Cozzi B, López-Albors O. 2022. The Heart of the Killer Whale: Description of a Plastinated Specimen and Review of the Available Literature. Animals (Basel) [Internet] 12. Available from: 10.3390/ani12030347

Li W, Yin L, Shen C, Hu K, Ge J, Sun A. 2018. Variants: Association With Cardiac Disorders. Front. Physiol. 9:1372.

Mass AM, Supin AY. 2018. Vision. In: Encyclopedia of Marine Mammals. Elsevier. p. 1035–1044.

McGowen MR, Grossman LI, Wildman DE. 2012. Dolphin genome provides evidence for adaptive evolution of nervous system genes and a molecular rate slowdown. Proc. Biol. Sci. 279:3643–3651.

McGowen MR, Montgomery SH, Clark C, Gatesy J. 2011. Phylogeny and adaptive evolution of the brain-development gene microcephalin (MCPH1) in cetaceans. BMC Evol. Biol. 11:98.

McGowen MR, Tsagkogeorga G, Álvarez-Carretero S, dos Reis M, Struebig M, Deaville R, Jepson PD, Jarman S, Polanowski A, Morin PA, et al. 2019. Phylogenomic Resolution of the Cetacean Tree of Life Using Target Sequence Capture. Syst. Biol. 69:479–501.

McGowen MR, Tsagkogeorga G, Williamson J, Morin PA, Rossiter ASJ. 2020. Positive Selection and Inactivation in the Vision and Hearing Genes of Cetaceans. Mol. Biol. Evol. 37:2069–2083.

Nery MF, Arroyo JI, Opazo JC. 2013. Genomic organization and differential signature of positive selection in the alpha and beta globin gene clusters in two cetacean species. Genome Biol. Evol. 5:2359–2367.

Nery MF, Arroyo JI, Opazo JC. 2014. Increased rate of hair keratin gene loss in the cetacean lineage. BMC Genomics 15:869.

Nery MF, González DJ, Opazo JC. 2013. How to Make a Dolphin: Molecular Signature of Positive Selection in Cetacean Genome. PLoS One 8:e65491.

Norris DP. 2012. Cilia, calcium and the basis of left-right asymmetry. BMC Biol. 10:102.

Nowak MA, Boerlijst MC, Cooke J, Smith JM. 1997. Evolution of genetic redundancy. Nature 388:167–171.

Oka K, Yamakawa M, Kawamura Y, Kutsukake N, Miura K. 2023. The Naked Mole-Rat as a Model for Healthy Aging. Annu Rev Anim Biosci 11:207–226.

Osipova E, Barsacchi R, Brown T, Sadanandan K, Gaede AH, Monte A, Jarrells J, Moebius C, Pippel M, Altshuler DL, et al. 2023. Loss of a gluconeogenic muscle enzyme contributed to adaptive metabolic traits in hummingbirds. Science 379:185–190.

Park JY, An Y-R, Kanda N, An C-M, An HS, Kang J-H, Kim EM, An D-H, Jung H, Joung M, et al. 2015. Cetaceans evolution: insights from the genome sequences of common minke whales. BMC Genomics 16:13.

Peichl L, Behrmann G, Kröger RH. 2001. For whales and seals the ocean is not blue: a visual pigment loss in marine mammals. Eur. J. Neurosci. 13:1520–1528.

Pennekamp P, Karcher C, Fischer A, Schweickert A, Skryabin B, Horst J, Blum M, Dworniczak B. 2002. The ion channel polycystin-2 is required for left-right axis determination in mice. Curr. Biol. 12:938–943.

Pinto B, Valente R, Caramelo F, Ruivo R, Castro LFC. 2023. Decay of Skin-Specific Gene Modules in Pangolins. J. Mol. Evol. [Internet]. Available from: 10.1007/s00239-023-10118-z

Poole K. 2022. The Diverse Physiological Functions of Mechanically Activated Ion Channels in Mammals. Annu. Rev. Physiol. 84:307–329.

Racicot R. 2022. Evolution of whale sensory ecology: Frontiers in nondestructive anatomical investigations. Anat. Rec. 305:736–752.

Randall JG, Gatesy J, Springer MS. 2022. Molecular evolutionary analyses of tooth genes support sequential loss of enamel and teeth in baleen whales (Mysticeti). Mol. Phylogenet. Evol. 171:107463.

Ridgway SH. 1988. The cetacean central nervous system. In: Comparative Neuroscience and Neurobiology. Boston, MA: Birkhäuser Boston. p. 20–25.

Rivaud MR, Delmar M, Remme CA. 2020. Heritable arrhythmia syndromes associated with abnormal cardiac sodium channel function: ionic and non-ionic mechanisms. Cardiovasc. Res. 116:1557–1570.

Sayers EW, Bolton EE, Brister JR, Canese K, Chan J, Comeau DC, Connor R, Funk K, Kelly C, Kim S, et al. 2022. Database resources of the national center for biotechnology information. Nucleic Acids Res. 50:D20–D26.

Scholander PF. 1940. Experimental Investigations on the Respiratory Function in Diving Mammals and Birds.

Schrider DR, Hahn MW. 2010. Gene copy-number polymorphism in nature. Proc. Biol. Sci. 277:3213–3221.

Shen H, Liu D, Wu K, Lei J, Yan N. 2019. Structures of human Na1.7 channel in complex with auxiliary subunits and animal toxins. Science 363:1303–1308.

Silva FA, Souza ÉMS, Ramos E, Freitas L, Nery MF. 2023. The molecular evolution of genes previously associated with large sizes reveals possible pathways to cetacean gigantism. Sci. Rep. 13:67.

Sleiman Y, Lacampagne A, Meli AC. 2021. “Ryanopathies” and RyR2 dysfunctions: can we further decipher them using in vitro human disease models? Cell Death Dis. 12:1041.

Sunami A, Glaaser IW, Fozzard HA. 2000. A critical residue for isoform difference in tetrodotoxin affinity is a molecular determinant of the external access path for local anesthetics in the cardiac sodium channel. Proc. Natl. Acad. Sci. U. S. A. 97:2326– 2331.

Sun X, Zhang Z, Sun Y, Li J, Xu S, Yang G. 2017. Comparative genomics analyses of alpha-keratins reveal insights into evolutionary adaptation of marine mammals. Front. Zool. 14:41.

Sun Y-B, Zhou W-P, Liu H-Q, Irwin DM, Shen Y-Y, Zhang Y-P. 2013. Genome-wide scans for candidate genes involved in the aquatic adaptation of dolphins. Genome Biol. Evol. 5:130–139.

Suyama M, Torrents D, Bork P. 2006. PAL2NAL: robust conversion of protein sequence alignments into the corresponding codon alignments. Nucleic Acids Res. 34:W609– W612.

Tarpley RJ, Hillmann DJ, Henk WG, George JC. 1997. Observations on the external morphology and vasculature of a fetal heart of the bowhead whale, Balaena mysticetus. Anat. Rec. 247:556–581.

Tatsuno R, Shikina M, Shirai Y, Wang J, Soyano K, Nishihara GN, Takatani T, Arakawa O. 2013. Change in the transfer profile of orally administered tetrodotoxin to non-toxic cultured pufferfish Takifugu rubripes depending of its development stage. Toxicon 65:76–80.

Tejada-Martinez D, de Magalhães JP, Opazo JC. 2021. Positive selection and gene duplications in tumour suppressor genes reveal clues about how cetaceans resist cancer. Proc. Biol. Sci. 288:20202592.

Thienel M, Müller-Reif JB, Zhang Z, Ehreiser V, Huth J, Shchurovska K, Kilani B, Schweizer L, Geyer PE, Zwiebel M, et al. 2023. Immobility-associated thromboprotection is conserved across mammalian species from bear to human. Science 380:178–187.

Tollis M, Ferris E, Campbell MS, Harris VK, Rupp SM, Harrison TM, Kiso WK, Schmitt DL, Garner MM, Aktipis CA, et al. 2021. Elephant Genomes Reveal Accelerated Evolution in Mechanisms Underlying Disease Defenses. Mol. Biol. Evol. 38:3606– 3620.

Tollis M, Robbins J, Webb AE, Kuderna LFK, Caulin AF, Garcia JD, Bèrubè M, Pourmand N, Marques-Bonet T, O’Connell MJ, et al. 2019. Return to the Sea, Get Huge, Beat Cancer: An Analysis of Cetacean Genomes Including an Assembly for the Humpback Whale (Megaptera novaeangliae). Mol. Biol. Evol. 36:1746–1763.

Turakhia Y, Chen HI, Marcovitz A, Bejerano G. 2020. A fully-automated method discovers loss of mouse-lethal and human-monogenic disease genes in 58 mammals. Nucleic Acids Res. 48:e91.

Uhlen M, Oksvold P, Fagerberg L, Lundberg E, Jonasson K, Forsberg M, Zwahlen M, Kampf C, Wester K, Hober S, et al. 2010. Towards a knowledge-based Human Protein Atlas. Nat. Biotechnol. 28:1248–1250.

Upham NS, Esselstyn JA, Jetz W. 2019. Inferring the mammal tree: Species-level sets of phylogenies for questions in ecology, evolution, and conservation. PLoS Biol. 17:e3000494.

Varró A, Tomek J, Nagy N, Virág L, Passini E, Rodriguez B, Baczkó I. 2021. Cardiac transmembrane ion channels and action potentials: cellular physiology and arrhythmogenic behavior. Physiol. Rev. 101:1083–1176.

Walker JR, Novick PA, Parsons WH, McGregor M, Zablocki J, Pande VS, Du Bois J. 2012. Marked difference in saxitoxin and tetrodotoxin affinity for the human nociceptive voltage-gated sodium channel (Nav1.7) [corrected]. Proc. Natl. Acad. Sci. U. S. A. 109:18102–18107.

Wang J, Chitsaz F, Derbyshire MK, Gonzales NR, Gwadz M, Lu S, Marchler GH, Song JS, Thanki N, Yamashita RA, et al. 2023. The conserved domain database in 2023. Nucleic Acids Res. 51:D384–D388.

Waterhouse A, Bertoni M, Bienert S, Studer G, Tauriello G, Gumienny R, Heer FT, de Beer TAP, Rempfer C, Bordoli L, et al. 2018. SWISS-MODEL: homology modelling of protein structures and complexes. Nucleic Acids Res. 46:W296–W303.

Williams TM, Fuiman LA, Kendall T, Berry P, Richter B, Noren SR, Thometz N, Shattock MJ, Farrell E, Stamper AM, et al. 2015. Exercise at depth alters bradycardia and incidence of cardiac anomalies in deep-diving marine mammals. Nat. Commun. 6:6055.

Würsig B, Thewissen JGM, Kovacs KM. 2017. Encyclopedia of Marine Mammals. Elsevier Science

Xie Z, Bailey A, Kuleshov MV, Clarke DJB, Evangelista JE, Jenkins SL, Lachmann A, Wojciechowicz ML, Kropiwnicki E, Jagodnik KM, et al. 2021. Gene Set Knowledge Discovery with Enrichr. Curr Protoc 1:e90.

Yang M, Derbyshire MK, Yamashita RA, Marchler-Bauer A. 2020. NCBI’s Conserved Domain Database and Tools for Protein Domain Analysis. Curr. Protoc. Bioinformatics 69:e90.

Yang Z. 2007. PAML 4: phylogenetic analysis by maximum likelihood. Mol. Biol. Evol. 24:1586–1591.

Yates AD, Allen J, Amode RM, Azov AG, Barba M, Becerra A, Bhai J, Campbell LI, Carbajo Martinez M, Chakiachvili M, et al. 2022. Ensembl Genomes 2022: an expanding genome resource for non-vertebrates. Nucleic Acids Res. 50:D996– D1003.

Yuasa T, Venugopal B, Weremowicz S, Morton CC, Guo L, Zhou J. 2002. The sequence, expression, and chromosomal localization of a novel polycystic kidney disease 1-like gene, PKD1L1, in human. Genomics 79:376–386.

Zhang X, Zong J, Chen S, Li M, Lu Y, Wang R, Xu H. 2020. Accumulation and Elimination of Tetrodotoxin in the Pufferfish by Dietary Administration of the Wild Toxic Gastropod. Toxins [Internet] 12. Available from: 10.3390/toxins12050278

Zhao Y, Seluanov A, Gorbunova V. 2021. Revelations About Aging and Disease from Unconventional Vertebrate Model Organisms. Annu. Rev. Genet. 55:135–159.

Zheng Z, Hua R, Xu G, Yang H, Shi P. 2022. Gene losses may contribute to subterranean adaptations in naked mole-rat and blind mole-rat. BMC Biol. 20:44.

